# Overlap graph-based generation of haplotigs for diploids and polyploids

**DOI:** 10.1101/378356

**Authors:** Jasmijn A. Baaijens, Alexander Schönhuth

**Affiliations:** Centrum Wiskunde & Informatica, Amsterdam, Netherlands; Utrecht University, Utrecht, Netherlands

## Abstract

Haplotype aware genome assembly plays an important role in genetics, medicine, and various other disciplines, yet generation of haplotype-resolved de novo assemblies remains a major challenge. Beyond distinguishing between errors and true sequential variants, one needs to assign the true variants to the different genome copies. Recent work has pointed out that the enormous quantities of traditional NGS read data have been greatly underexploited in terms of haplotig computation so far, which reflects that methodology for reference independent haplotig computation has not yet reached maturity. We present POLYTE (POLYploid genome fitTEr) as a new approach to de novo generation of haplotigs for diploid and polyploid genomes. Our method follows an iterative scheme where in each iteration reads or contigs are joined, based on their interplay in terms of an underlying haplotype-aware overlap graph. Along the iterations, contigs grow while preserving their haplotype identity. Benchmarking experiments on both real and simulated data demonstrate that POLYTE establishes new standards in terms of error-free reconstruction of haplotype-specific sequence. As a consequence, POLYTE outperforms state-of-the-art approaches in various relevant aspects, where advantages become particularly distinct in polyploid settings. POLYTE is freely available as part of the HaploConduct package at https://github.com/HaploConduct/HaploConduct, implemented in Python and C++.

## 1 Introduction

In most eukaryotic organisms genomes come in copies, where each copy can be assigned to one of the ancestors. The number of copies determines the *ploidy* of the organism where *diploid* relates to two copies, and *polyploid* refers to more than two copies^1^. The copies generally differ, because they carry their own sets of genetic variants; the copy-specific sequences are referred to as *haplotypes.* Distinguishing the two (in diploid organisms, such as in most vertebrates) or more than two (in polyploid organisms, such as many plants and some funghi) haplotypes plays an important role in various disciplines. Prominent examples are genetics, where assigning variants to ancestors is key (Tewhey *et al.*, 2014), and medicine, because very often haplotype-specific combinations of variants establish clinically relevant effects, for example, when disease risks have been inherited (Glusman *et al.*, 2014). In general, determining *haplotypic sequence* and thereby keeping track of ancestry based dependencies is instrumental in many settings.

Assembling the two (diploid) or more (polyploid) haplotypes from sequencing reads is known as *haplotype aware genome assembly*, and the resulting assembled pieces of sequence are referred to as *haplotigs.* The advent of next-generation sequencing (NGS) has resulted in a plethora of NGS read compatible assembly programs. However, the vast majority of these programs yield consensus genome sequence, as a summary across all haplotypes involved, instead of haplotigs. Even this is affected by fundamental challenges, because NGS reads contain errors and are relatively short. Often also hardware limitations can pose obstacles during the assembly process that are hard to deal with.

Generating haplotigs from NGS reads—which is the challenge that we tackle here—comes with additional obstacles. Beyond distinguishing between errors and true sequential variants, one needs to assign the true sequential variants to the different copies. This requires keeping track of information that allows to link the true sequential variants stemming from identical copies. However, NGS reads are rather short in general: techniques are needed that can link haplotype-specific variants across read boundaries. This is not necessarily a standard procedure in genome assembly, despite the many recent advances. In summary, haplotype aware assembly can still be considered in its early stages of development; every advance is desirable.

### Motivation

The majority of sequencing machines installed worldwide perform traditional NGS, such as Illumina sequencing. A plethora of population-scale sequencing studies (e.g. (Besenbacher *et al.*, 2015; Sudmant *et al.*, 2015; The Genome of the Netherlands Consortium, 2014; The UK10K Consortium, 2015)) have filled up databases with traditional, short NGS reads. In terms of quantities, traditional short NGS reads exceed the amount of reads stemming from more recent third-generation-sequencing (TGS) protocols by orders of magnitude. TGS protocols have considerably spurred the development of methods for haplotype-aware assembly (see Related work), because of the increase in read length. However, their increase in sequencing error rates is also a disturbing factor when dis tinguishing between haplotypes, which leaves applicants with ambiguities.

Recent work has pointed out that targeted examination of NGS (Illumina type) reads can have significant positive effects in haplotype aware assembly (Berger *et al.*, 2014; Patterson *et al.*, 2015). Seemingly, the enormous quantities of traditional NGS read data have been underexploited in terms of haplotig computation so far. This establishes our major motivation.

To better understand where serious progress can be made, one needs to realize that existing methods for haplotype computation from traditional NGS (Illumina) reads fall into two classes: the first (and arguably more popular) choice of approaches are referred to as *haplotype assembly* programs. These approaches make use of a reference genome to call variants from aligned reads, which are subsequently phased into separate haplotypes. The advantage of haplotype assembly programs is their stability and their resource-friendly usage. Examples for *diploid haplotype assembly* are WhatsHap (Patterson *et al.*, 2015), Phaser (Castel *et al.*, 2016), Hap-Cut2 (Edge *et al.*, 2017), ProbHap (Kuleshov, 2014) and HapCol (Pirola *et al.*, 2016). Examples for *polyploid haplotype assembly* are Hap-Compass (Aguiar and Istrail, 2012), HapTree (Berger *et al.*, 2014), SDhaP (Das and Vikalo, 2015), and H-PoP (Xie *et al.*, 2016). The disadvantage of haplotype assembly programs is that they depend on high-quality reference sequence as a backbone, and, in addition, also on external variant call sets, which are major external factors that can introduce non-negligible biases.

The second class of methods are *de novo haplotype aware assembly* approaches that can deal with traditional NGS (in particular Illumina) reads. The advantage of such approaches is that they are independent of reference genomes and external call sets, which eliminates the externally induced biases. There are only little such approaches available however; to the best of our knowledge, only ALLPATHS-LG (Ribeiro *et al.*, 2012), Platanus (Kajitani *et al.*, 2014), and dipSPAdes (Safonova *et al.*, 2015) explicitly aim at computation of haplotigs from (diploid) NGS data. However, ALLPATHS-LG and Platanus require particularly tailored libraries, which renders their general application difficult, and the dipSPAdes software is no longer maintained. In results of ours, we further noted that SPAdes (Bankevich *et al.*, 2012) can be run in diploid mode (which is not to be confused with the no longer maintained dipSPAdes), and is able to compute haplotigs (surprisingly not only in diploid, but also in conventional mode), thereby likely establishing the only tool among the (myriad of) approaches for consensus oriented genome assembly (see Bradnam *et al.* (2013); Salzberg *et al.* (2011) for references).

In summary, there are no approaches that 1) specialize in the generation of (high-quality) haplotigs, but 2) do not depend on high quality reference sequence as a backbone, 3) do not depend on external variant call sets and 4) do not require particularly tailored sequencing libraries.

### Contribution

The contribution of this paper is to close this gap in the landscape of approaches. We present POLYTE (POLYploid genome fitTEr), as an approach to do this. Our results indicate that POLYTE outperforms state-of-the-art approaches of the two classes, where advantages are significant in a variety of relevant aspects. As an example of an application scenario, POLYTE outperforms the other approaches in reconstructing individual haplotypes of the MHC region. This is a critical application because of the high genetic variability of this region, which renders its reconstruction particularly challenging. Note finally that the majority of approaches focuses on diploid genomes. Therefore, the lack of approaches that can compute haplotigs for polyploid organisms is even more striking. POLYTE achieves performance rates for settings of higher ploidy that are nearly on a par with those achieved for diploid organisms. To the best of our understanding, one might perceive POLYTE’s achievements for polyploid organisms as a novelty in its own right.

### Related work

In terms of assembly paradigms, POLYTE is an *overlap graph based* approach. It follows an iterative scheme, where in each iteration, an overlap graph is constructed, where nodes represent reads (in the initial step) or contigs (in the subsequent steps). Edges indicate that a pair of reads/contigs is likely to stem from identical haplotypes. Based on their interplay with other reads or contigs in terms of the underlying overlap graph, reads or contigs are joined and subsequently transformed into new contigs, which then become the nodes of the overlap graph of the following iteration. Thereby, contigs grow along the iterations, while preserving their haplotype identity. POLYTE adopts ideas from earlier work that either focused on variant discovery (Marschall *et al.*, 2012), viral quasispecies assembly (Baaijens *et al.*, 2017; Töpfer *et al.*, 2014) or metagenome gene assembly (Gregor *et al.*, 2016). However, Marschall *et al.* (2012) only make use of a rudimentary type of (overlap like) graph, where edges are agnostic to sequence content, and does not follow an iterative scheme. Second, Gregor *et al.* (2016) deal with protein sequence, rather than DNA. Finally, Baaijens *et al.* (2017) and Töpfer *et al.* (2014) focus on *deep coverage settings* and require at least a few 100x per haplotype. In this respect, POLYTE unites the virtues of Marschall *et al.* (2012)—the ability to handle low coverage—on the one hand, and Baaijens *et al.* (2017); Töpfer *et al*. (2014) —dealing with real overlap graphs and contig computation—on the other hand. That is, POLYTE brings forward an iterative overlap graph based scheme for contig generation that reliably works in *low coverage settings*, requiring coverage of only as low as 5x per haplotype.

Note finally that our approach also draws motivation from the recent technology shifts, such as the advent of third-generation sequencing (TGS) and explicitly haplotype-aware sequencing protocols like StrandSeq (Porubsky *et al.*, 2017), which have put the computation of haplotigs into the focus of current attention. Chin *et al.* (2016); Jain *et al.* (2018); Weisenfeld *et al.* (2017) describe approaches that aim to exploit the respective advances in sequencing technology and protocol design. We consider the adaptation of POLYTE to TGS data most interesting future work: the framework of POLYTE is generic in terms of choosing reads, such that this is a matter of adapting parameters, more than anything else. We recall, however, that our motivation was to bring forward a method that exploits (the abundantly available) traditional NGS reads in the first place. This, for example, enables to reconstruct MHC region haplotypes in various population-scale studies (e.g. Besenbacher *et al.* (2015); Sudmant *et al.* (2015); The Genome of the Netherlands Consortium (2014); The UK10K Consortium (2015)), which has been a major challenge so far.

## 2 Methods

We present POLYTE, an algorithm to assemble individual haplotypes of diploid and polyploid genomes from short read sequencing data; see Figure 1 for the complete workflow. POLYTE follows the overlap-layout-consensus (OLC) paradigm, where consensus refers to removing errors within haplotypes (instead of the common interpretation of reaching consensus across different haplotypes). Our method starts by constructing a *read overlap graph* which is used for error correction of the input sequences. Subsequently, we make use of an iterative OLC scheme, where in each iteration a *contig overlap graph* is constructed. This graph is further reduced by applying *transitive edge removal* and *read-based branch reduction*. Then, contigs are clustered and merged according to their interplay within the overlap graph, resulting in a collection of extended contigs (‘contig extension’ in Figure 1). These extended contigs establish the nodes of the contig overlap graph of the next iteration, which is achieved by an updating procedure. When contigs can not be merged any further, POLYTE outputs the final set of contigs. When dealing with diploid organisms, an additional assembly stage can be activated which consists of two additional steps (‘diploid branch reduction’ and ‘contig extension’ in Figure 1), creating an optional output that is refined for diploid organisms.

**Figure 1:**
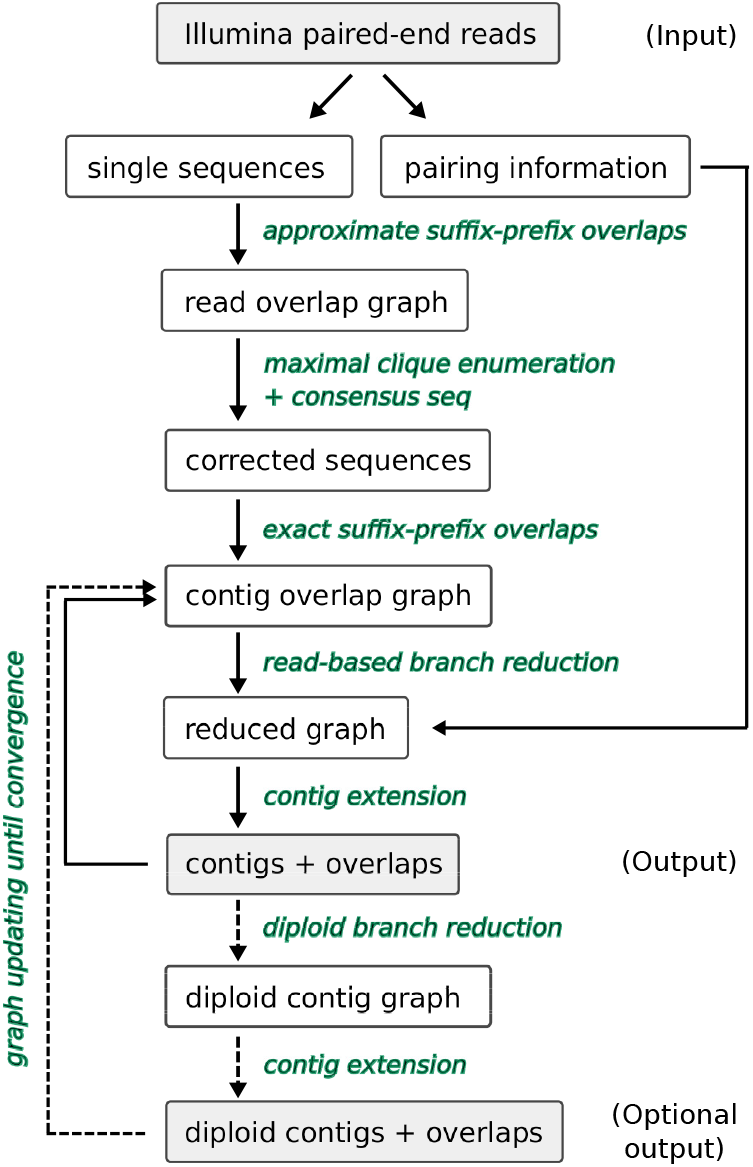
Algorithm overview.

Given that we are dealing with data of relatively low sequencing depth, we need to exploit the information present in the sequencing reads as much as possible. The initial error correction procedure is particularly crucial, as sequencing errors can heavily disturb the process of distinguishing between different haplotypes. For this error correction step, approximate suffix-prefix overlaps are computed to establish an initial read overlap graph. Inspired by Baaijens *et al.* (2017) and Töpfer *et al.* (2014), maximal cliques are enumerated and errors are corrected by inspecting the read overlaps within the cliques. By design of the overlap graph—edges indicate that two reads stem from identical haplotypes—every clique only contains reads from identical haplotypes, which allows to eliminate errors based on majority votes. Note that this procedure is particularly tailored to low coverage settings with known ploidy: admissible clique sizes and minimal sequence overlap lengths can heavily vary in comparison to earlier approaches. However, with edge criteria that are much less restrictive than in other approaches, we obtain a larger number of spurious edges. We have developed a procedure for *read-based branch reduction* to reduce the number of spurious edges in the overlap graph, which is of great importance for accurate reconstruction of haplotigs.

In the following sections we will discuss each of the steps involved in POLYTE, following the workflow depicted in Figure 1.

### 2.1 Read overlap graph construction

The steps outlined in this section refer to the initial step ‘approximate suffix-prefix overlaps’ that leads to the establishment of the ‘read overlap graph’ in Figure 1.

#### Read overlap graph: definition

The read overlap graph follows the idea that nodes are reads and edges indicate that a pair of reads stem from identical haplotypes. Given the input consisting of paired-end sequencing reads (Il-lumina), let 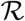 be the collection of single end sequences from all paired-end reads. The *read overlap graph G = (V, E)* is a directed graph where *V* corresponds to the collection of input sequences 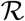. That is, for every paired-end read we have two vertices *υ,υ′* ∈ *V*, one for each single end sequence 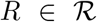. Directed edges *υ_i_* → *υ_j_* ∈ *E* connect sequences *R_i_, R_j_* whenever the suffix of *R_i_* overlaps the prefix of *R_j_* for at least 50% of the average sequence length. Furthermore, for each edge *υ_i_ → υ_j_*, we require *QS*(*R_i_, R_j_*) ≥ *δ*, where 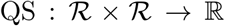 is a quality score and *δ* is an appropriate threshold, determined based on empirical statistics so as to maximize the chances that the edge (*υ_i_,υ_j_*) in deed indicates that the corresponding sequences *R_i_* and *R_j_* stem from identical haplotypes. In this, we largely follow ideas presented in earlier work (Baaijens *et al.*, 2017; Marschall *et al.*, 2012; Töpfer *et al.*, 2014). The difference with respect to these prior approaches is that only single ends are considered, whereas in the earlier approaches nodes represent the entire paired-end reads. Also note that here overlap graphs are twice as large in comparison to the earlier approaches, because each paired-end read is represented by two nodes, instead of only one. While this difference imposes substantial methodical and technical challenges, it is key to dealing with low coverage because it decisively increases the recall in terms of recovering reads that stem from identical haplotypes.However, it also implies follow-up complications, because the information that read ends come in pairs is temporarily lost. In POLYTE, paired-end information is stored and used in later steps; see Section 2.4 below.

#### Construction

Computation of the edges for the read overlap graph requires enumeration of all pairwise approximate suffix-prefix overlaps (of sufficient length) between the single read ends 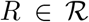 and evaluation of a quality score *QS*(*R_i_, R_j_*) for each pair of sequences for which a sufficiently good overlap was established during the approximate suffix-prefix overlap computation. We further orient the edges (which is necessary because reads can stem from either the forward or the reverse stand) and systematically remove double transitive edges, which ensures that one can enumerate maximal cliques in an efficient manner (see Section 2.2). Each of these graph construction steps is described in detail in Section 1 of the Supplementary Material.

The computation of approximate suffix-prefix overlaps for vertebrate genome sized input read sets is a serious issue, currently hardly conceivable without external auxiliary means (see also Simpson and Durbin (2012)). Here, we suggest a method that aims to suppress externally introduced biases to a maximum degree. We make use of a reference genome for binning reads in an initial step and, after binning, we discard the reference genome and any related information entirely such that POLYTE operates in full de novo mode.

### 2.2 Correction of sequencing errors

After the establishment of the read overlap graph, we cluster its nodes by enumerating the *maximal cliques* contained in it. The idea is to collect groups of reads belonging to the same haplotype and produce error-free sequences for subsequent assembly steps (‘corrected sequences’, Figure 1). By definition of a maximal clique—a maximal group of nodes all of which are connected by edges—maximal cliques represent maximum-sized groups of reads all of which belong to the same haplotype. Once all maximal cliques are determined, it is therefore reasonable to merge the reads within a maximal clique into a single contig. Note that this contig is longer than the individual reads participating in the contig and that sequencing errors can be eliminated by raising majority votes among the reads participating in the maximal clique. While this reflects an approved procedure in its generic form (Baaijens *et al.*, 2017; Gregor *et al.*, 2016; Töpfer *et al.*, 2014), accounting for the particular setting we are facing here—namely low coverage in combination with sequence-based edge definition—requires particular care.

We make the minimum clique size depend on the coverage per haplotype; in all settings considered we are dealing with known ploidy, while the overall coverage of reads can be determined by usual considerations, which yields per-haplotype-coverage estimates. To determine the optimal minimum size of a clique for a given per-haplotype coverage, we compute the probability *p_c,k_* that, due to unfortunate fragmentation of sequencing reads, there is no clique of size *k* that extends a given sequencing read R to the right when requiring at least 50% read overlap. In other words, we compute the probability that there are *at most k*–1 reads extending *R* to the right; the exact same analysis applies to extensions to the left.

For determining *p_c,k_*, we assume that sequencing reads are fragmented randomly, which implies that reads are generated independently of one another. Let *R* be a read and *S* be a set containing reads from the haplotype of *R* at exactly 1x coverage, further assuming that all reads *R*′ ∈ *S* have the same length as R (which reflects that all single read ends have the same length). It is straightforward to see that the probability that there is *R′ ∈ S* that overlaps *R* at at least 50% of its length (into one direction, left or right) as 0.5. When dealing with a per-haplotype coverage of *c*, we assume the existence of *c* sets of reads *S_i_,i* = 1,…,*c* all of which contain reads that cover the haplotype *c* at 1x. For computing *p_c,k_* we consider that for only *k* – 2 of the *c* sets *S_i_, i* = 1,…, *c* we have that there is *R*′ ∈ *S_i_* that overlaps *R* at at least 50% of its length (resulting in a clique of size at most *k* – 1), which evaluates as

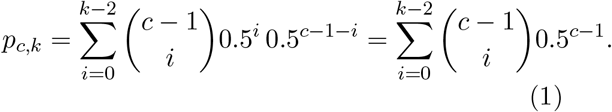

We aim to have *p_c,k_* low to be able to deal with sufficiently many cliques, hence for every choice of *c* we compute k such that *p_c,k_* < 0.001. In this regard, we obtain that for up to 10x per haplotype an appropriate choice for the minimum clique size is 2, for coverages between 10x and 15x a minimum clique size of 3 is required, while for *c* ≥ 15x an optimal choice for the minimum clique size is 4. Note that cliques cannot grow larger than size 4 because of double transitive edge removal (see Supplementary Material).

### 2.3 Contig overlap graph construction

Given the corrected sequences obtained by merging maximal cliques, we build a new graph: the contig overlap graph (see Figure 1).

#### Contig overlap graph: definition

The contig overlap graph *G*′ = (*V′,E′*) is very similar to the read overlap graph, except that we construct it from a set of contigs assumed to be free of sequencing errors. Therefore, every node *υ ∈ V′* corresponds to a contig and we add an edge between a pair of nodes whenever they have an exact (i.e. error-free) overlap of sufficient length.

#### Construction

The contig overlap graph can be constructed very efficiently by making use of the FM-index-based algorithm from Section 2.1 while allowing only exact overlaps. This gives us the complete edge set *E*′ without any further computations, since we do not need to compute the overlap quality score for exact overlaps. Note that the minimal overlap length in the contig overlap graph does not need to be as high as before error correction and it is independent of the read length: all experiments were performed using a minimal contig overlap of 50bp.

### 2.4 Branch reduction in the contig overlap graph

Before using the contig overlap graph to extend our contigs, we trim the graph by removing redundant vertices and edges and resolving branches based on read evidence where possible, now also exploiting the paired-end information. After completing this step, we have a ‘reduced graph’ (see Figure 1) that is ready for contig extension.

#### Transitive edge removal

An edge *u* → *w ∈ E′* is called *transitive* if there exists a vertex *υ ∈ V′* and edges *u → υ, υ → w ∈ E′*. Now that sequences (contigs) are assumed to be error-free, transitive edges have become fully redundant, hence we remove all transitive edges from the graph before further processing.

#### Branching edges and nodes

The *indegree* (resp. *outdegree*) of a node *υ ∈ V′* is defined as the total number of incoming (resp. outgoing) edges in *G*″. If *υ* has indegree greater than one, we say v has an *in-branch*; analogously, if v has outdegree greater than one, we say that *υ* has an *out-branch*. We refer to the corresponding edges as *branching edges* and to *υ* as a *branching node*. Since we did not use any read pairing information during construction of our overlap graphs, we observe many branches in the contig overlap graph. We now use the information how ends are paired to remove any branching edges in the contig overlap graph that do not correspond to a true haplotype.

#### Merging simple paths

Following the above definition, any edges that are not branching edges constitute *simple paths* through the contig overlap graph. For such paths, there is only one possible way to combine the corresponding contigs; hence, before processing the graph any further, we merge every simple path into a single contig. Since edges in the graph represent exact overlaps, this is a straightforward procedure.

#### Branching components

After merging simple paths, all remaining edges are branching edges. We define a *branching component* as subgraph H of the contig overlap graph, such that (1) *H* is an induced subgraph, (2) *H* is connected as an undirected graph, and (3) within *H*, any vertex has only incoming or outgoing edges in *H*, but not both. A branching component is defined to be maximal with respect to these three properties; see Figure 2. Intuitively, a branching component reflects all possible haplotypes within a small region of the genome.

**Figure 2:**
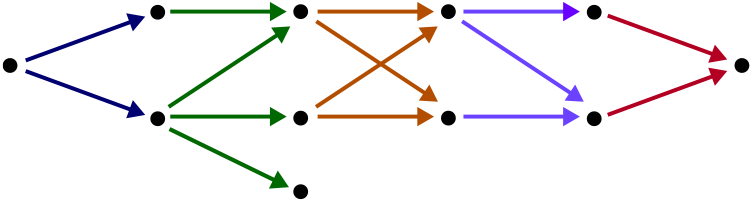
Illustration of branching components in a contig overlap graph. Edges of the same color belong to the same branching component.

Note that different components may intersect across their vertex sets, but cannot have any edges in common. In other words, the maximal branching components partition the set of all branching edges, as illustrated in Figure 2. This partition can be found in time linear in the number of branching edges by alternatingly traversing in-branch edges and out-branch edges until every edge has been seen exactly once; see Section 2 in the Supplementary Material for further details. After enumerating all maximal branching components, we evaluate read evidence per component.

#### Read evidence

The main idea of read-based branch reduction is to remove all branching edges for which there is insufficient *read evidence* in the input data. For this purpose, we keep track of all original sequencing reads (‘subreads’) that were used to build a contig; each of these subreads may provide evidence for a branching edge. Within a branching component, we first list all *variant positions*, i.e., the positions at which the sequences corresponding to the different neighbors differ from each other. Intuitively, these are the positions where we may find sequencing reads supporting a given branching edge. A paired-end sequencing read *R* = (*R*_1_, *R*_2_) is marked as *evidence* for the branching edge *u* → *υ* if it satisfies the following conditions:

i. *R* spans the branching edge, meaning that at least one of the sequences *R*_1_, *R*_2_ is a subread of *u and* at least one of the sequences *R*_1_, *R*_2_ is a subread of *υ*;
ii. The sequence spanning the edge is identical to the contig sequence of the corresponding node for all variant positions it covers;
iii. *R* is unique for this edge: it does not satisfy conditions (i) and (ii) for any other edge involved in this branching component.

Figure 3 shows two examples of contigs creating branches in the overlap graph, along with the sequencing reads (‘subreads’) that were used to build these contigs; the subreads providing read evidence are highlighted in yellow. Observe that in panel A, in order to satisfy condition (iii) a subread has to cover at least one variant position on either contig. In panel B, we illustrate that also a single read end can provide evidence: the rightmost subread covers a variant position and satisfies all conditions listed above.

**Figure 3:**
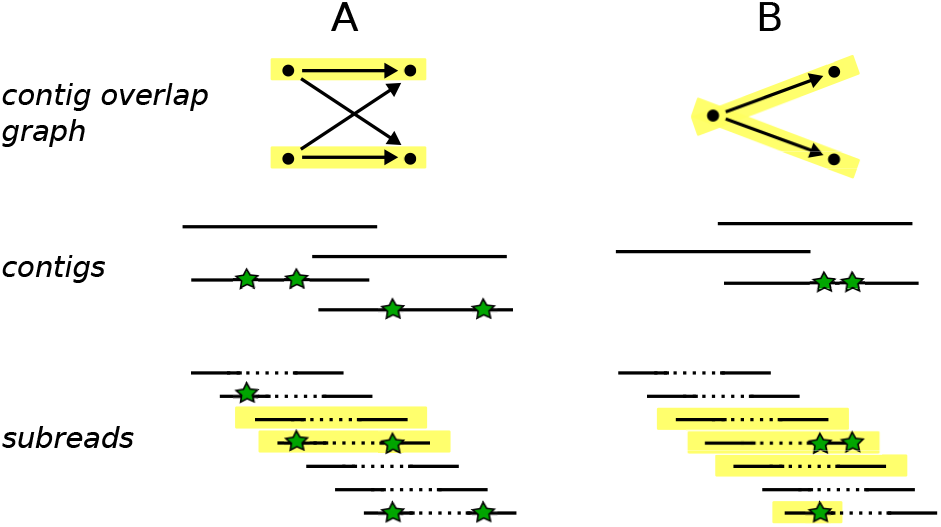
Two examples of contigs creating branches in the overlap graph. Edges corresponding to true haplotypes are highlighted in yellow. The corresponding subreads are aligned below, those providing read evidence are again highlighted.

Note that condition (ii) ensures that erroneous contigs do not find evidence in correct reads: if a sequencing error accidentally ends up in a contig, it will cause a branch in the overlap graph which can only be supported by reads containing exactly this sequencing error. Whenever such a branch occurs, there will be insufficient evidence and hence the erroneous contigs will never be merged. Eventually, these contigs can be filtered out based on their short length. In the Supplementary Material (Section 2) we discuss how an appropriate evidence threshold is determined (using similar considerations as for determining the optimal clique size, Section 2.2).

#### Branching edge removal

For every branching component, we count the read evidence per branching edge and remove any edges with evidence count below the evidence threshold.

### 2.5 Contig extension and graph updating

After applying the read-based branch reduction techniques described above, all branches have been either resolved or removed from the contig overlap graph. Contig extension has become an easy task: any contigs which are connected by an edge in the graph must belong to the same haplotype, and, therefore, we merge each such pair of contigs into a new, longer contig. Then, we update the overlap graph: the extended contigs become the new nodes and the edges are updated accordingly. The resulting updated graph is used for further assembly in an iterative manner, as described in Section 2.6.

### 2.6 Iterative procedure and diploid mode

Our workflow consists of iteratively performing the steps described in Sections 2.3–2.5, as illustrated in Figure 1. The algorithm terminates when the edge set *E*′ of the updated contig overlap graph becomes empty, either upon construction or after branch reduction. Thus, our algorithm is guaranteed to converge, and once it does we remove any remaining inclusions from the final contig set. Also any contigs shorter than the fragment size of the original reads are removed from the output.

#### Diploid mode

Knowing that a given sample is diploid is a very strong piece of information when performing haplotype assembly. We have developed a special module which can be activated for diploid samples. It extends the POLYTE pipeline by two additional steps after the standard algorithm has terminated: construction of a diploid contig graph, followed by contig extension (see Figure 1). In these additional steps, we use the knowledge that the sample is diploid to resolve additional branches (for which there was insufficient evidence in the read set to resolve them during the read-based branch reduction step; see Section 2.4).

In overlap graphs from diploid samples we typically see two types of branching components; Figure 4 illustrates both types (Panel A and B) and gives an example of a possible collection of contigs giving rise to the corresponding branching component. In both situations we have four contigs, two from each haplotype, which have identical sequence where the contigs overlap. In diploid mode, a single read of evidence may already be considered sufficient, depending on the amount of evidence found for the other edges (Supplementary Material, Section 3).

**Figure 4:**
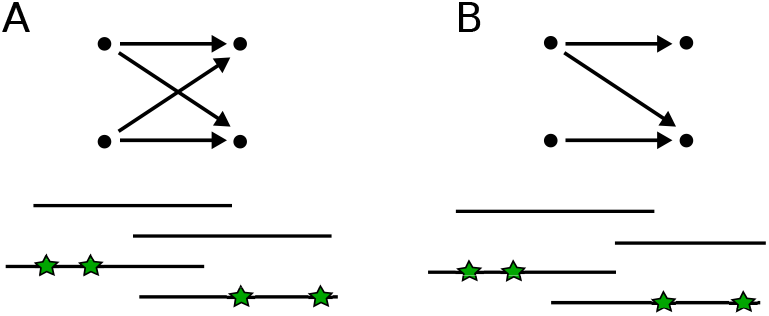
Typical branching components in diploid assemblies: four contigs, two from each haplotype, having identical sequence in their overlap. Depending on the contig lengths, all contigs overlap (panel A) or only a subset of the contigs overlap (panel B).

This procedure is more risky than default branch reduction (Section 2.4), since it does not require such stringent read evidence. Therefore, we always run the main POLYTE algorithm until convergence before turning to diploid mode (Figure 1). This ensures that all evidence in the original reads has been exploited first.

## 3 Results

In this section we show results for POLYTE on both simulated and real Illumina data sets and evaluate the assembly quality in terms of haplotype coverage, N50, NGA50, error rate, and misassembled contig length relative to the total assembly length. We also compare our method against alternative haplotype reconstruction tools: SPAdes (Bankevich *et al.*, 2012), Phaser (Castel *et al.*, 2016), HapCut2 (Edge *et al.*, 2017), WhatsHap (Patterson *et al.*, 2015), SGA (Simpson and Durbin, 2012), and H-PoP (Xie *et al.*, 2016). Other polyploid assemblers (Aguiar and Istrail, 2012; Berger *et al.*, 2014; Das and Vikalo, 2015) were unable to process our benchmarking data. All methods were run with default settings and assembly statistics were obtained with QUAST (Gurevich *et al.*, 2013).

### 3.1 Data sets

#### Simulated data

We generated a collection of simulated data sets of varying ploidy and sequencing depth to evaluate the effect of these characteristics. We selected four human MHC haplotypes from the Vega Genome Browser^2^: COX, DBB, MANN, and SSTO. Subsequently, we used SimSeq^3^ to simulate Illumina MiSeq reads of length 2×250bp for each of those haplotypes at a coverage of 5x, 10x, 20x, 30x, 40x, and 50x, respectively, and combined the resulting read sets to form data sets of ploidy 1 (only COX haplotype, a sanity check), ploidy 2 (COX and DBB), ploidy 3 (COX, DBB, and MANN), and ploidy 4 (all).

#### Real data

For evaluation on real sequencing data, we considered a data set from phase 3 of the 1000 Genomes project (1000 Genomes Project Consortium *et al.*, 2012; Sudmant *et al.*, 2015) for individuals NA19240. This data set was obtained from a 2×250 bp PCR free Illumina protocol, sequenced to a coverage of 28-68x. Full haplotypes have been reconstructed for this individual as part of a recent study (Chaisson *et al.*, 2017) using various specialized sequencing techniques and reconstruction algorithms; we use the resulting haplotypes as a ground truth for a whole-chromosome benchmarking experiment on chromosome 22.

#### Alignments and variant call sets for reference-guided methods

Reference-guided methods Phaser, HapCut2, WhatsHap, and H-PoP require as input a reference genome, read alignments to the reference genome, and a precomputed set of genomic variants. For the simulated data we performed read alignment to the GRCh38 reference genome using BWA MEM (Li and Durbin, 2009). The real data was already provided as alignments to the GRCh37 reference genome, also obtained with BWA MEM. We extracted the sequencing reads corresponding to chromosome 22 from the provided BAM files. Finally, we performed variant calling on all data sets with FreeBayes^4^.

### 3.2 Assembly performance criteria

We evaluate assembly performance in terms of several statistics commonly used for de novo assembly evaluation, as reported by QUAST.

#### Haplotype coverage (HC)

The completeness of the assembly is measured by the fraction of nucleotides in the target haplotypes (ground truth) covered by haplotigs, referred to as the haplotype coverage.

#### N50 and NGA50

Assembly contiguity is measured using the N50 value, which is defined as the length for which the collection of all contigs of that length or longer covers at least half the assembly. The NGA50 measure is computed in a similar fashion, but only aligned blocks are considered (obtained by breaking contigs at misassembly events and removing all unaligned bases). This measure reports the length for which the total size of all aligned blocks of this length or longer equals at least 50% of the total length of the true haplotypes.

#### Error rate (ER) and N-rate (NR)

We evaluate error rate as the sum of mismatch rate and indel rate when comparing to the ground truth haplotype sequences. In addition, we report the relative number of ambiguous bases (‘N’s), referred to as the N-rate.

#### Misassembled contig length (MCL)

A contig or haplotig is called misassembled if it contains at least one misassembly, meaning that left and right flanking sequences align to the true haplotypes with a gap or overlap of more than 1kbp, or align to different strands, or even align to different haplotypes. The misassembled contig length is defined as the total number of bases in misassembled contigs relative to the total assembly length.

### 3.3 Benchmarking results

We performed benchmarking experiments on one of the simulated MHC data sets described above (ploidy 2, 20x coverage per haplotype) to compare a variety of haplotype reconstruction tools. In addition, we ran all methods on the chromosome 22 data of the 1000 Genomes individual NA19240. The assembly statistics on both data sets are shown in Table 1. Since both data sets are diploid, we present results for SPAdes in regular mode and in diploid mode, referred to as SPAdes-dip.

**Table 1:** Benchmarking results, HC = Haplotype Coverage, ER = Error Rate (mismatches + indels), NR = N-Rate (ambiguous bases), MCL = Misassembled Contig Length. Top: simulated diploid data for the MHC region. Bottom: real data for chromosome 22 of 1000 Genomes individual NA19240. HC values within parentheses indicate haplotype coverage relative to the amount of bases covered by sequencing reads.

|  | HC (%) | N50 | NGA50 | ER (%) | NR (%) | MCL (%) |
| --- | --- | --- | --- | --- | --- | --- |
| *Simulated data* |  |  |  |  |  |  |
| POLYTE | 92.4 | 4397 | 4394 | 0.035 | 0 | 0 |
| SGA | 73.4 | 3444 | - | 0.025 | 0 | 0 |
| SPAdes | 84.1 | 3588 | 919 | 0.032 | 0 | 0.8 |
| SPAdes-dip | 83.6 | 3294 | 903 | 0.003 | 0 | 0 |
| HapCut2 | 84.5 | 29259 | 17980 | 0.068 | 0 | 7.1 |
| H-PoP | 81.7 | 32319 | 17484 | 0.158 | 0 | 12.7 |
| Phaser | 82.6 | 24785 | 16884 | 0.095 | 0 | 5.1 |
| WhatsHap | 85.2 | 32656 | 17980 | 0.098 | 0 | 10.6 |
| *Real data* |  |  |  |  |  |  |
| POLYTE | 78.2 (90.5) | 2838 | 2316 | 0.090 | 0 | 0.4 |
| SGA | 57.7 (66.8) | 2842 | - | 0.069 | 0 | 0.0 |
| SPAdes | 67.0 (77.5) | 5798 | - | 0.131 | 0 | 1.7 |
| SPAdes-dip | 66.4 (76.9) | 5772 | - | 0.139 | 0 | 2.3 |
| HapCut2 | 70.1 (81.1) | 6541 | 5306 | 0.090 | 0.9 | 1.0 |
| H-PoP | 62.4 (72.2) | 9583 | 7435 | 0.119 | 0.9 | 1.9 |
| Phaser | 66.2 (76.6) | 6394 | 5245 | 0.094 | 0.9 | 0.8 |
| WhatsHap | 67.6 (78.2) | 6257 | 6094 | 0.092 | 0.9 | 0.7 |

In both experiments, we observe that across all methods POLYTE has the largest haplotype coverage (HC, 92.4% and 78.2% for MHC and chr22, respectively). In other words, it reconstructs the largest fraction of the true haplotype sequences. In comparison, the other methods are all more or less on a par (81.7–85.2% [MHC] and 57.7–70.1% [Chr22], respectively). On the real data the haplotype coverage achieved by all methods is rather low; this can be explained by only 86.4% of the target haplotypes being covered by sequencing reads. After normalizing the haplotype coverage values by 86.4, POLYTE achieves a haplotype coverage of 90.5%.

In terms of assembly contiguity, indicated by high N50 and NGA50 values, reference-guided methods (HapCut2, Phaser, WhatsHap, H-PoP) perform better than de novo assemblers (POLYTE, SGA, SPAdes). This reflects a common advantage of reference-guided approaches, which can make use of the external information to bridge regions only poorly covered with informative reads, if appropriate. The increase in length, however, is offset by a substantial decrease in terms of haplotig quality: reference-guided approaches exhibit both substantially more misassemblies (which in particular can lead to severe issues in downsteam interpretations) and increased error rates, here larger by one to two orders of magnitude. Note that several NGA50 values are undefined (‘-’), because the aligned blocks are unable to cover at least 50% of the total reference length.

Another important difference between reference-guided methods and de novo approaches is reflected in the N-rates on the real data: the reference genome contains several stretches of ambiguous nucleotides (‘N’s), which the reference-guided methods cannot correct. De novo approaches, on the other hand, can potentially uncover the true sequence behind these ambiguous regions and show an N-rate of 0% (versus 0.9% for the reference-guided methods).

Between de novo approaches, we compare POLYTE with SGA and SPAdes and observe that POLYTE reconstructs a substantially larger fraction of the true haplotypes. Although SPAdes achieves better N50 values, this comes at the expense of a decrease in terms of error rate and misassemblies, also reflected in a low NGA50 value on the simulated data and the NGA50 being undefined on the real data (see explanation above).

On the simulated data set, POLYTE and SPAdes achieve comparable error rates of 0.035% and 0.031%, respectively. On the real data we notice an advantage for POLYTE, with an error rate of only 0.090% compared to 0.131% for SPAdes. In addition, POLYTE is less vulnerable to misassemblies than SPAdes, with 0% versus 0.8% MCL on simulated data and 0.4% versus

1. 7% MCL on real data. SGA is able to reconstruct highly accurate contigs with slightly lower error rates than POLYTE (0.025 vs 0.035% [MHC] and 0.069 vs 0.090% [Chr22], respectively), but covers a significantly lower fraction of the ground truth haplotypes (73.4 vs 92.4% [MHC] and 57.7 vs 78.2% [Chr22], respectively).

In an overall account, we believe that, arguably, the major advantage of POLYTE is established by the increase of 10-15% over the other approaches in terms of haplotype-specific coverage, in combination with the error rates, which are clearly lower than those of the other tools.

In terms of runtime and memory usage, de novo approaches are in general more expensive than reference-guided methods. We also observe this when comparing CPU time and peak memory usage (Supplementary Tables 1 and 2). Reference-guided methods have CPU times that are orders of magnitude less compared to de novo methods (where POLYTE requires 9–15 times more (resp. 3–6 times more) runtime and 3 times less (resp. 12–17 times more) memory than SPAdes and SGA, respectively). It is important to notice, however, that these de novo assemblers are highly parallelizable, while reference-guided methods are not. This leads to feasible runtimes on multi-core computing facilities in practice.

### 3.4 Effect of ploidy and sequencing depth

To study the effect of genome ploidy and sequencing depth on the assembly quality and completeness, we ran POLYTE, SPAdes, SGA, and H-PoP on all simulated data sets described in Section 3.1 (other tools were unsuitable for polyploid genomes). Figure 5 shows the results for the 5x, 10x, and 20x data sets in terms of haplotype coverage (HC), N50, NGA50, error rate (ER), and misassembled contig length (MCL). For additional result tables we refer the reader to the Supplementary Tables 3–6.

**Figure 5:**
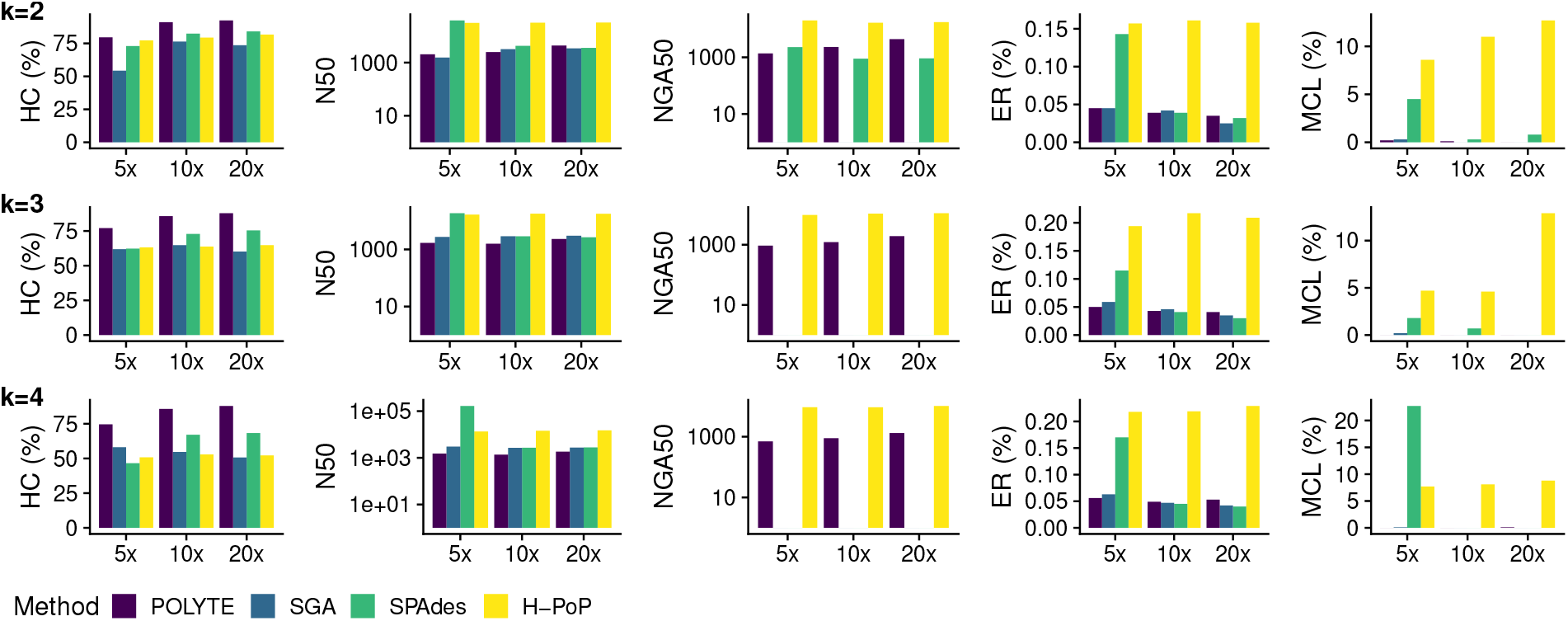
Assembly results per method for simulated data of increasing ploidy (k=2,3,4) and per-haplotype coverage (5x,10x,20x). N50 and NGA50 values are plotted on a log-scale for increased readability. For SGA (k=2,3,4) and SPAdes (k=3,4) the NGA50 values are undefined.

We observe that POLYTE excels regarding haplotype coverage, with advantages becoming more distinct as the ploidy increases. SPAdes and H-PoP achieve more contiguous assemblies (higher N50 values) but, as we already observed on diploid data, this comes at the cost of significantly higher error rates and misassemblies. SGA performs very similar to POLYTE when considering N50, ER, and MCL, but obtains much lower HC values. The NGA50 values highlight the improved assembly quality of POLYTE over SGA and SPAdes: while POLYTE achieves NGA50 values comparable to the N50, SGA and SPAdes are unable to cover at least 50% of the ground truth with alignments (hence NGA50 is undefined). Overall, we conclude that in polyploid settings the same advantages of POLYTE apply as in diploid settings - in creased haplotype-specific coverage in combination with low error rates – and become even more pronounced.

All other methods evaluated (HapCut2, Phaser, Whatshap, SPAdes-dip) are designed for diploid data, so for those we could only assess the effect of sequencing depth. Results indicate that each of the reference-guided methods already performs optimally at a coverage per haplotype of 5x. Moreover, these methods are unaffected by a further increase in sequencing depth (see Supplementary Table 7). SPAdes in diploid mode (SPAdes-dip) performs optimally at a per-haplotype coverage 20x.

## 4 Discussion

Assembling the individual haplotypes of an organism from sequencing reads is known as *haplotype aware genome assembly* and plays a major role in various disciplines, including genetics and medicine (Glusman *et al.*, 2014; Tewhey *et al.*, 2014). Computing haplotype-specific pieces of sequence, also known as *haplotigs*, is a difficult task. Algorithms addressing this task do not only need to distinguish between sequencing errors and true variants, but also need to assign the true variants to the individual haplotypes. Enormous quantities of next-generation sequencing (NGS) reads generated worldwide have not been fully exploited in terms of haplotig computation, because methodology for de novo haplotig computation from NGS reads has been in a rather immature state.

We have presented POLYTE (POLYploid genome fitTEr) as a new approach to de novo assembly of haplotigs from NGS data, suitable for diploid genomes as well as genomes of higher ploidy. Unlike the majority of NGS based de novo assemblers, our method follows the overlap-layout-consensus (OLC) paradigm to achieve enhanced performance rates in terms of haplotype-specific computation of contigs. In order to appropriately distinguish between errors and true variants to be assigned to haplotypes, it employs an iterative OLC scheme. Along the iterations, contigs grow in length while preserving their uniqueness in terms of haplotype identity. As a result, POLYTE outperforms the currently available state-of-the-art approaches for haplotig computation, where it performs particularly favorable in terms of quantities that refer to haplotype-specific reconstruction of the genomes. We showed that POLYTE succeeds in accurate reconstruction of individual haplotypes of the human MHC region, including the highly polymorphic HLA genes. Future work may therefore be to apply POLYTE to NGS data in population-scale human genome projects, where individual genomes are still lacking proper annotation of their MHC region, which applies in the majority of cases. Advantages become particularly distinct on data of higher ploidy, leading to plant genome assembly as another interesting future application of POLYTE.

Our algorithm is, in its essence, generic in the choice of input reads, so applying it for TGS reads essentially is a matter of adapting parameters, which we will explore in the short-term future.

## Funding

This work was supported by the Netherlands Organisation for Scientific Research (NWO) through Vidi grant 679.072.309.

## Footnotes

1 Depending on the context, polyploid includes diploid, but here we refer to polyploid as more than two copies.

2 http://vega.archive.ensembl.org/info/data/MHC_Homo_sapiens.html

3 https://github.com/jstjohn/SimSeq

4 https://github.com/ekg/freebayes

